# A Locus Control Region Generates Distinct Active Placental Lactogen And Inactive Growth Hormone Gene Domains In Term Placenta That Are Disrupted With Obesity

**DOI:** 10.1101/2024.12.18.628357

**Authors:** Yan Jin, Ian McNicol, Peter A. Cattini

## Abstract

A Placental villi include an outer layer of syncytiotrophoblasts (STBs) and an inner layer of cytotrophoblasts (CTBs) that fuse to generate STBs in pregnancy. While activation of the single locus containing the human (h) placental lactogen (hPL) genes (*hPL-A*/*CSH1* and *hPL-B*/*CSH2*) begins in the CTBs, their expression in STBs requires further epigenetic modifications as well as interactions between locus control region (LCR) and gene regulatory sequences. No transcription factor that limits or facilitates hPL LCR/gene interactions for locus activation is reported but the paternally-expressed gene 3 (PEG3/PW1) transcription factor is a candidate. PEG3 is expressed by villous CTBs but not STBs, and putative PEG3 sites were identified in the hPL LCR and promoter sequences. Furthermore, dysregulation of both hPL and PEG3 gene expression have been linked to peripartum depression. Using CTB-like JEG-3 cells, we show PEG3 binding to hypersensitive sites (HS III-V) within the LCR, and that hPL transcript levels increase with PEG-3 knockdown. In term placenta, PEG3 binding at placenta-specific HS IV was increased with maternal obesity, where a decrease in hPL RNA levels is seen, while PEG3 binding was reduced in women with obesity who develop insulin-treated gestational *diabetes mellitus* (O/GDM+Ins), where increased hPL gene expression is observed. Chromatin conformation capture revealed distinct hPL gene domain interactions that are modified with maternal obesity but largely reversed in O/GDM+Ins, correlating with PEG3 binding. Thus, decreased PEG3 binding may be required for hPL domain generation and expression during CTB to STB transition.

## Introduction

Placental villi include an outer layer of syncytiotrophoblasts (STBs) and an inner layer of cytotrophoblasts (CTBs) that can grow and fuse to generate and maintain the STB layer throughout human (h) pregnancy (Mayhew 2014; Mori, et al. 2007). The human (h) placental lactogen (PL), also known as chorionic somatomammotropin hormone (CSH), is a pregnancy-related metabolic adapter that is produced by the STBs (McWilliams and Boime 1980; Newbern and Freemark 2011). The hPL genes include *hPL-A*/*CSH1* and *hPL-B*/*CSH2* that code independently for the same PL, as well as the *hCS-L*/*CSHL1* pseudogene for which there is RNA but no known functional protein product (Giampietro, et al. 1984; Walker, et al. 1990). These genes are located together in tandem with those coding for pituitary growth hormone (*hGH-N*/*GH1*) and placental growth hormone (*hGH-V*/*GH2*) and, as products of gene duplication, all share greater than 90% nucleotide sequence homology (Bewley and Li 1984; Chen, et al. 1989). Activation of the hPL genes begins during the transition of villous CTBs to STBs, but further modifications and interaction between a remote locus control region (LCR) and local gene sequences separated by 35 kb is required for hPL gene expression by STBs (Jin, et al. 2018; Kimura, et al. 2007).

The LCR includes three hypersensitive sites (HS III-V) (Elefant, et al. 2000a; Jin, et al. 2004). HS III possesses enhancer activity and binds activator protein (AP)-2α (Jin et al. 2004), while HS IV, which is placenta-specific, and HS V are linked to locus boundary element activity through association with CCCTC binding factor (CTCF) (Elefant et al. 2000a; Jin, et al. 2011; Jin et al. 2004; Jones, et al. 1995; Tsai, et al. 2014). Regulatory regions near the hPL genes, such as upstream P sequences and downstream 3′-regulatory region (3′-RR), also play roles in modulating gene expression by interacting with multiple transcription factors (Chen et al. 1989; Elefant, et al. 2000b; Jacquemin, et al. 1994; Jiang, et al. 1999; Lytras, et al. 2011; Norquay, et al. 2006; Vakili, et al. 2013; Walker et al. 1990). However, no transcription factor that limits hPL locus activation and presumably LCR/gene interactions in the transitioning CTB is reported but the paternally-expressed gene 3 (PEG3/PW1) transcription factor is a candidate (Cattini, et al. 2020).

PEG3 is a zinc finger transcription factor expressed from the paternal allele that binds the consensus sequence 5′-gkGGswsT-3′ site with high affinity and regulates multiple cellular processes including potentially as a transcriptional repressor (Cattini et al. 2020; Kuroiwa, et al. 1996; Lee, et al. 2015; Sojoodi, et al. 2016; Thiaville, et al. 2013; Torres, et al. 2017). Putative PEG3 binding sites are present in both the hPL LCR and proximal promoter regions (Cattini et al. 2020). Furthermore, PEG3 is expressed in various villous CTB subtypes, including syncytin-2-positive CTBs that fuse to generate the multinucleated STB layer, but not by the STBs themselves (Hiby, et al. 2001; Liu, et al. 2018; Mayhew 2014; Mori et al. 2007). Together, these observations support a potential role for PEG3 limiting hPL gene locus activation and/or expression.

Here, we investigated PEG3’s ability to bind the hPL LCR and promoter sequences in CTB-like JEG-3 cells and term placentas, where variation in levels of hPL gene expression by maternal obesity and gestational *diabetes mellitus* (GDM) are reported (Jin et al. 2018). Our findings support a role for PEG3 in the regulation of hPL gene expression by repression and/or limiting activation of the hPL locus, which requires a specific chromosomal architecture that is disrupted with maternal obesity and associated with ongoing or increased PEG3 binding at HS IV. Schematic representations of these regulatory interactions are presented.

## Materials and Methods

### Tissue and cell samples

Placenta samples were obtained from three groups of women after approval of the Health Research Ethics Board at the University of Manitoba, and with signed informed consent forms from all participants. Term placentas were obtained from three groups: (1) lean women with a body mass index (BMI) 20-25; (2) women with obesity (BMI >35) but no gestational complications; and (3) women with obesity subsequently diagnosed with GDM and treated with insulin (O/GDM+Ins). The clinical characteristics of pregnancies of these three groups of women from whom term placenta samples were used here for quantitative real-time reverse transcription polymerase chain reaction (qPCR), chromatin conformation capture (3C) and chromatin immunoprecipitation (ChIP)-qPCR analyses were described previously (Jin et al. 2018). BMI was calculated from pre-pregnancy weight and height, and basic clinical characteristics of the pregnancies from which term placental samples were taken and used for analyses were previously reported (Jin et al. 2018). Human astrocytoma U87, embryonic kidney 293 (HEK293) and choriocarcinoma BeWo and JEG-3 cells were cultured as described (Ganguly, et al. 2015; Yang, et al. 2010).

### RNA isolation and quantitative real-time reverse transcription polymerase chain reaction (PCR)

Placental tissue samples (10 mm^3^) were collected from the chorion and processed to enrich for STBs (Jin et al. 2018; Vakili et al. 2013). RNA from tissues and cell lines was isolated using the RNeasy Plus mini kit (Qiagen), and quality was confirmed by agarose gel electrophoresis. Total RNA (1 μg) was converted to cDNA using the QuantiTect reverse transcription kit and quantitative PCR (qPCR) was performed using specific primers (Table 1). Specific amplifications were identified by a single peak in the melting curve and expression levels were calculated from standard curves for primer sets and normalized to GAPDH expression (Jin et al. 2018).

**Table 1.**
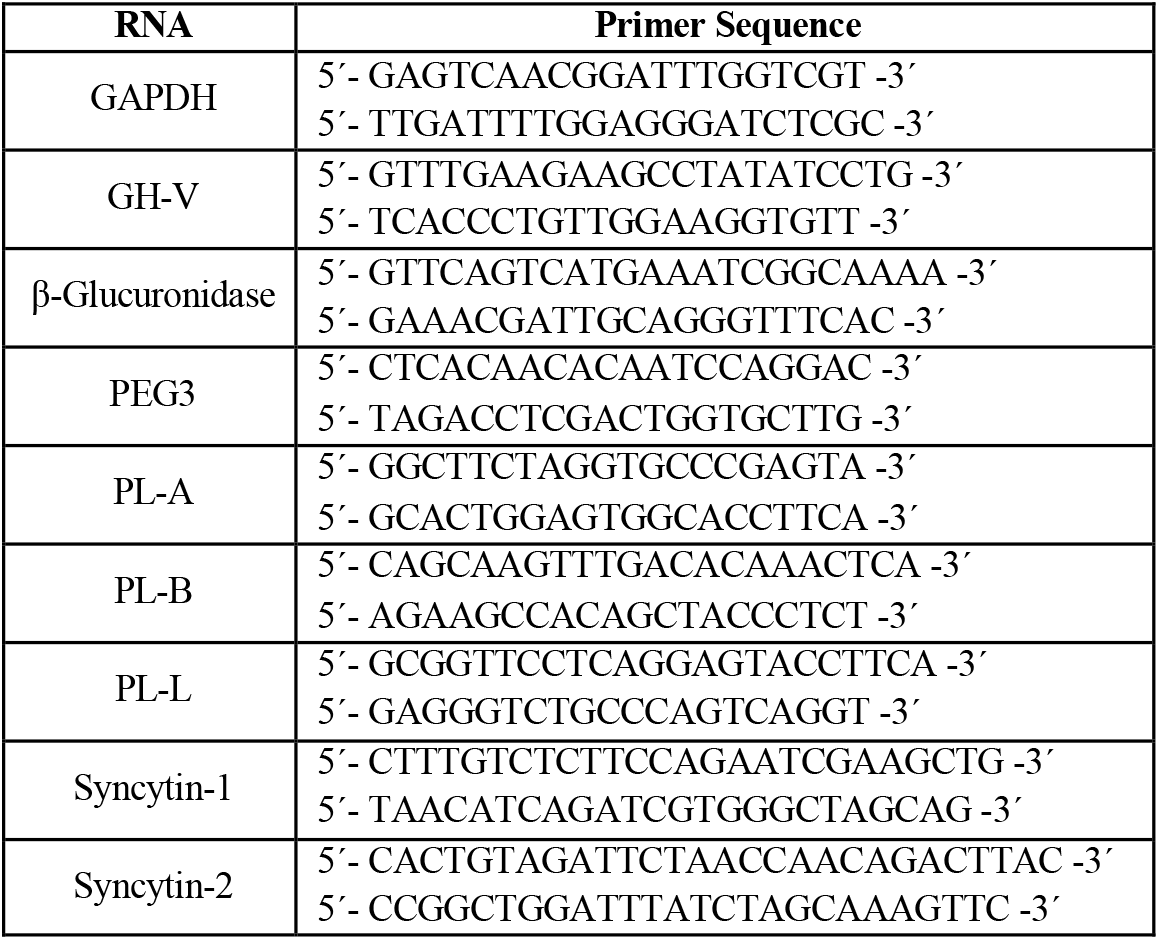
PCR primers used to detect human RNA levels.

### Lentiviral-mediated shRNA knockdown

Two PEG3-targeting shRNAs were obtained from Origene (https://www.origene.com). JEG-3 cells (1×10^5^) were seeded in 2 ml of RPMI and 10% v/v fetal bovine serum growth medium, transduced with 30 MOI of lentiviral particles, and polybrene (final concentration 8 μg/ml). After 24 hours, viral particles were removed and cells were cultured for 72 hours before RNA isolation.

### Electrophoretic mobility shift assay (EMSA)

Competition-EMSA was done as described previously (Jin, et al. 1999) with JEG-3 cell nuclear extract isolated using the EpiSeeker nuclear extraction kit (Abcam). Nuclear extract (3 μg), was pre-incubated with 50- or 100-fold molar excess of competitor oligonucleotides (Table 2) and 2 μg of poly(dI-dC) for 5 minutes, before adding ^32^P-radiolabeled phosphoglucomutase-2-Like-1 (Pgm2L1) oligonucleotide probe (1 ng) and the reactions were incubated for a further 20 minutes at 4 ^o^C. Complexes were separated on non-denaturing 5% (w/v) polyacrylamide gels and visualized by autoradiography.

**Table 2.**
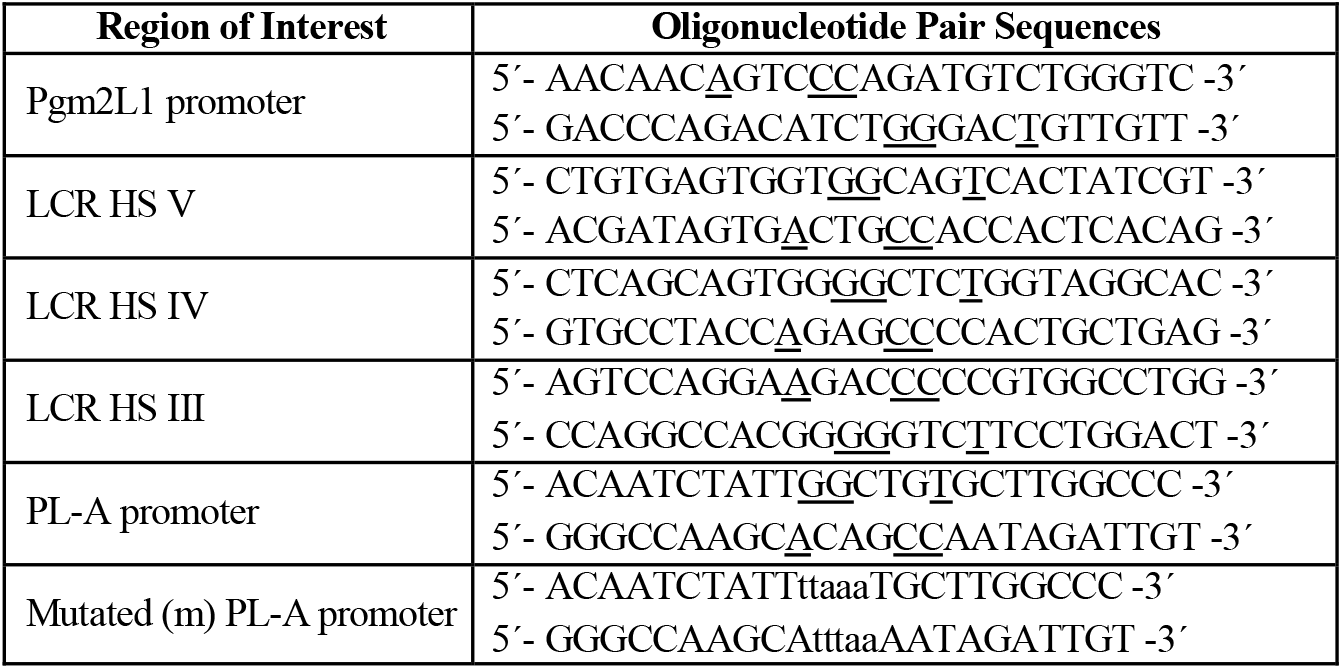
Oligonucleotide sequences used for EMSA containing a known or putative PEG3-like DNA element based on the presence of a GG-T motif (underlined) and match with a consensus PEG3 site (5′-g t/g G G c/g a/t g/c T-3′). The sequences modified in the PEG3-like site contained within the distal human PL-A promoter site, which are common to the PL-A, PL-B, and PL-L as well as GH-N and GH-V gene promoter sequences, are indicated by lower case letters.

### Chromatin immunoprecipitation (ChIP)

Chip-qPCR was performed by Active Motif (https://www.activemotif.com). Briefly, chromatin was extracted and sheared. ChIP DNA enriched from three independent assays were verified by qPCR using the human Myoferlin (MYOF), transcription enhancer factor (TEF)-1 and Glioma Tumor Suppressor Candidate Region 1 (GLTSCR1) 1 genes as positive controls, using a polyclonal anti-PEG3 antibody (Invitrogen PA5-55282), and a gene desert on human chromosome 12 (Untrl 12) as a negative control. Assays were done using 50 μg of chromatin and 30 μl of antibody in triplicate. ChIP DNA enrichment is graphed as percentage of input (the relative amount of immunoprecipitated DNA compared to 1% input DNA after qPCR analysis). The regions assessed and primers used are listed in Table 3.

**Table 3.**
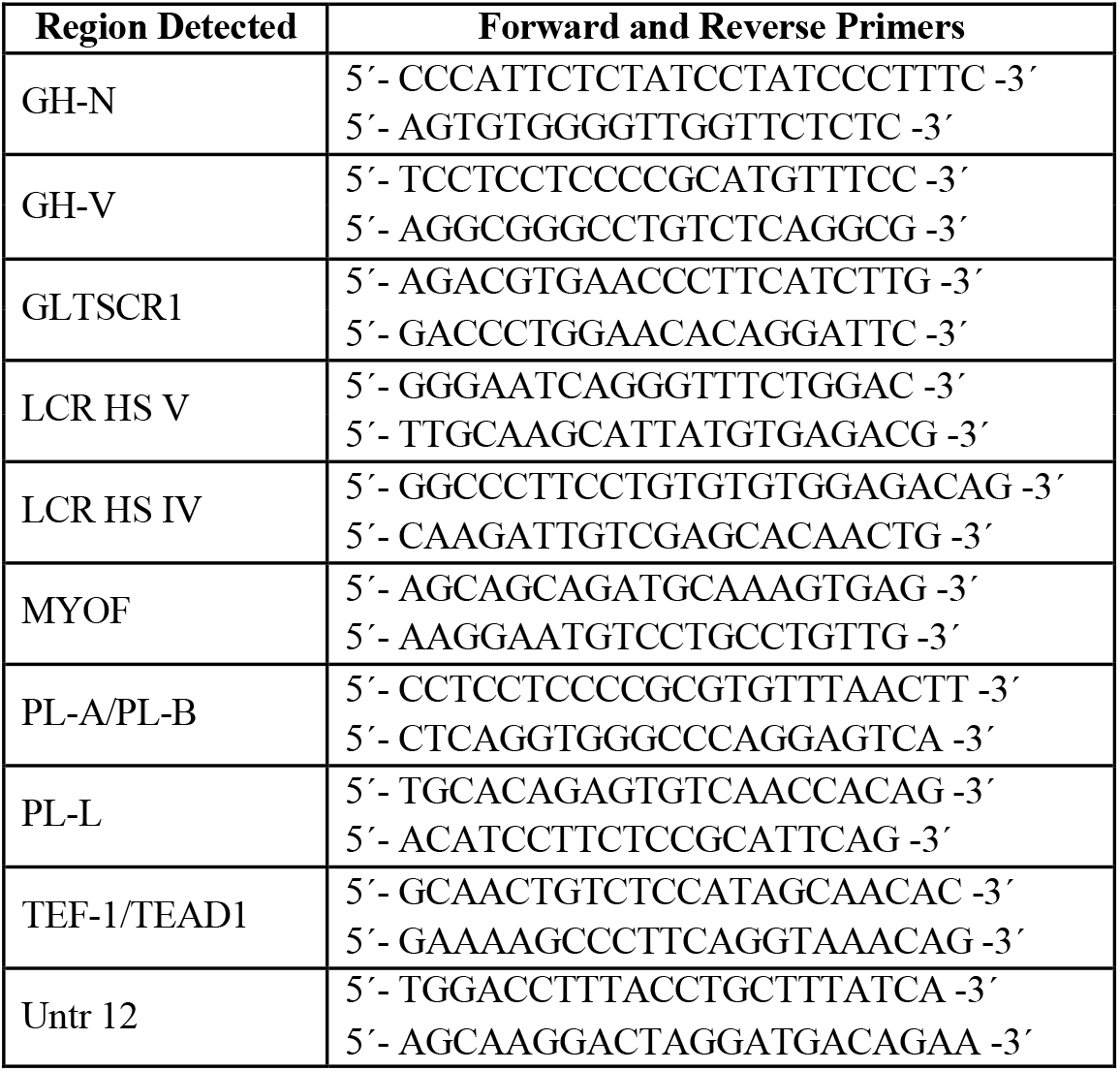
Primers used for ChIP-qPCR.

### Chromosome conformation capture (3C) assay

STB nuclei isolated from multiple regions of placenta chorion were used for 3C assay as described previously (Aliakbar, et al. 1990; Jin et al. 2018; Tsai, et al. 2016). Briefly, nuclei were cross-linked with 1% formaldehyde, digested with *Bgl*II, and ligated. Ligated DNA was reverse-crosslinked and purified. Two rounds of PCR with nested primers (Table 4) were done to detect interactions between HS III, P sequences and 3′-RRs. The efficiency of digestion was assessed by qPCR using a primer pair that encompassed the *Bgl*II cut site with the Power SYBR PCR Kit (#4368702; Applied Biosystems, MA, USA) (Jin et al. 2018). The results were normalized to hGAPDH in uncut and cut samples. Only samples with over 85% digestion were analyzed.

**Table 4.**
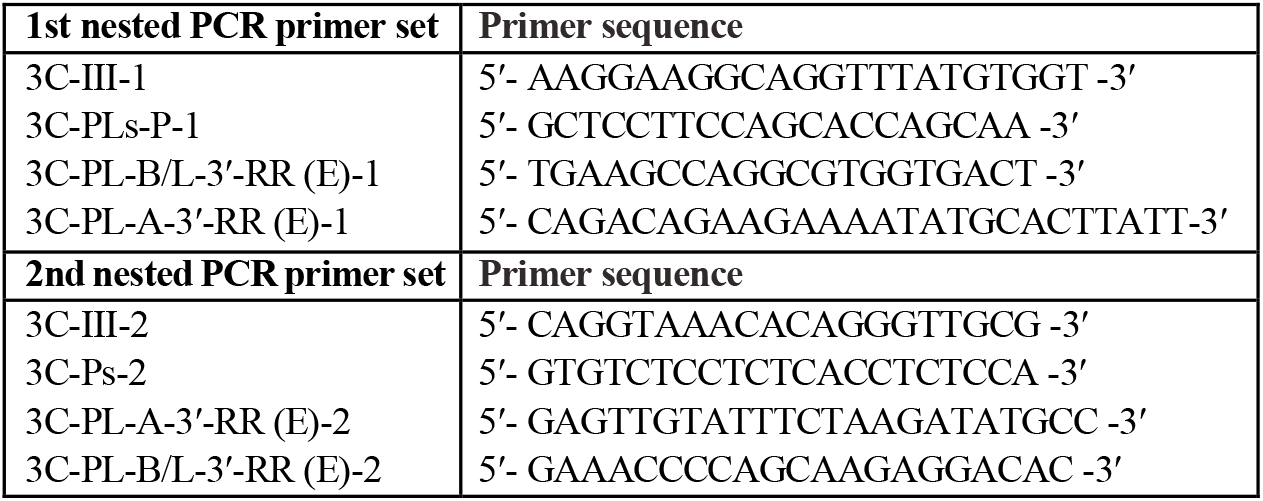
Primers used for 3C assay.

### Amplicon sequencing and bioinformatics

Amplicon sequencing was done by Applied Biological Materials Inc. (Richmond, BC, Canada) and the Biomedical Research Centre (Vancouver, BC, Canada) as described previously (Jin et al. 2018). Briefly, the PCR products from 3C assay ligation were purified, subjected to end-repair, adapter ligation, and PCR amplification to generate a library of amplicons for sequencing. Sequences were compared to our predicted 3C assay ligation products database and the numbers of sequences representing each interaction product were counted and the ratio of each interaction was calculated as percentage over the total numbers of matched sequences, which were >3000 for HS III, P sequence and 3′-RR interactions tested. The ligation frequency is presented as a percentage of the ligation frequency detected in the placentas of (lean) women with a BMI <25, which was arbitrarily set to 100%. All 3C assay ligation products were normalized to their corresponding internal control DNA (hGAPDH).

### Statistical analysis

Data were analyzed using GraphPad Prism 10. For single comparisons, a two-tailed t-test was applied. Multiple comparisons used one-way ANOVA or Kruskal-Wallis test with appropriate *post hoc* tests. A value of p<0.05 was considered statistically significant. Error bars represent the standard error of the mean.

## Results

Relative PEG3 RNA levels were assessed in CTB-like JEG-3 and BeWo cells as well as non-placental human glioblastoma U87 cells and embryonic kidney (HEK) 293 cells by qPCR (Figure 1A). PEG3 transcripts were detected in placental JEG-3 and BeWo as well as non-placental U87 cells but were low or undetectable in HEK293 cells. Relative PEG-3 RNA levels were significantly greater in JEG-3 cells when compared to either BeWo (∼3.6-fold, p<0.0001, n=6) or U87 (∼4.7-fold, p<0.0001) cells (Figure 1A).

**Figure 1.**
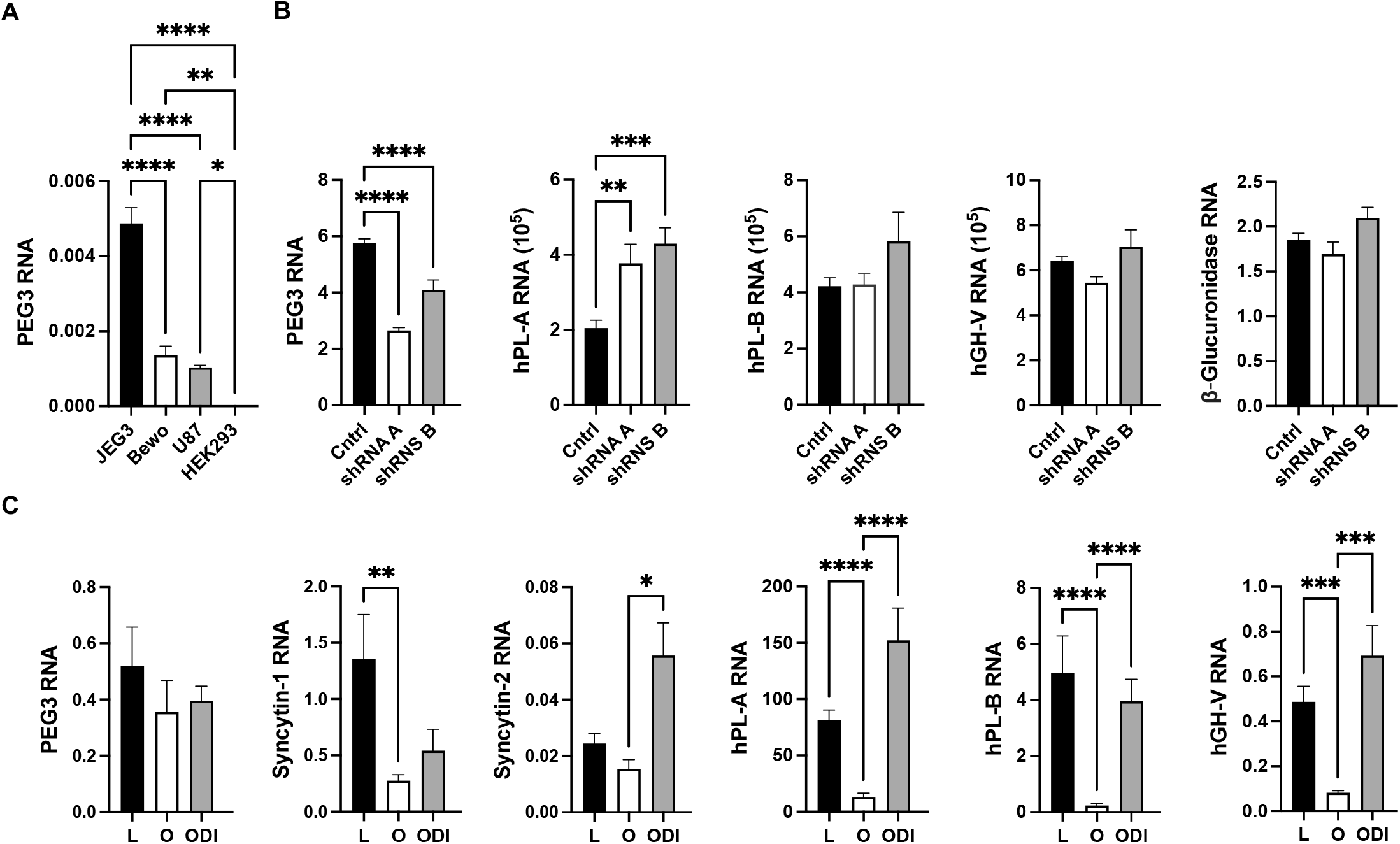
**(A)** PEG3 RNA levels relative to GAPDH transcripts were assessed in JEG-3, BeWo, U87 and HEK293 cells by qPCR, and calculated from a standard curve assuming similar reaction kinetics. Results were assessed by one-way ANOVA with Tukey’s *post hoc* test (n=8). **(B)** JEG-3 cells were treated with scrambled/control (Cntrl) or PEG3 shRNAs A or B. PEG3, hPL-A, hPL-B, hGH-V and glucuronidase (Gluc) RNA levels were assessed by qPCR 72 hours later. Results were analyzed by one-way ANOVA with Dunnet’s multiple comparison test (n=6). **(C)** Term placenta samples from lean women (L; BMI <25), women with obesity (O; BMI >35) and women with O/GDM+Ins (ODI) were assessed for PEG-3, syncytin-1 and syncytin-2 RNA levels by qPCR. Results were analyzed with the Kruskal-Wallis test (n=8-13). Results reported for hPL-A, hPL-B and minimal hGH-V RNA levels (Jin et al. 2018) are re-analyzed here for comparison. Values presented are the mean plus or minus standard error of the mean and significant differences are indicated by: ^*^ p<0.05; ^**^ p<0.01; ^***^ p<0.001; and ^****^ p<0.0001.

To assess a direct effect of reduced PEG3 availability, JEG3 cells were infected with lentivirus expressing one of two (A and B) short hairpin (sh) RNAs for PEG3 relative to a scrambled shRNA control for 72 hours. PEG-3, hPL-A, hPL-B, hGH-V and glucuronidase RNA levels were assessed by qPCR. No significant effects were detected at 10 MOI (not shown), however, a significant ∼40% decrease in PEG3 was detected with PEG3 shRNAs at MOI 30 (Figure 1B). A corresponding modest but significant ∼2-fold increase in hPL-A RNA levels was seen with both PEG3 shRNA A and B, however, an increase in hPL-B and hGH-V transcript levels were only suggested with PEG3 shRNA B but was not significant (Figure 1B). By contrast, no effects of PEG3 knockdown on glucuronidase transcript levels were observed with PEG3 shRNA A and B relative to a scrambled shRNA control (Figure 1B).

RNA levels for PEG3 as well as syncytin-1 and syncytin-2, both markers of human trophoblast cell fusion (Vargas, et al. 2009), were compared in term placenta samples from lean women, women with pre-pregnancy obesity and women with O/GDM+Ins by qPCR (Figure 1C). No significant effect of maternal obesity with or without GDM and insulin treatment was detected on PEG3 transcript levels when compared to placenta samples from lean women. This contrasts with the decrease in hPL/GH-V RNA levels seen with maternal obesity, and the reversal of this negative effect in women with GDM+Ins (Jin et al. 2018), which is reassessed here (Figure 1C). An ∼69% decrease in syncytin-1 RNA levels was observed with maternal obesity compared to lean women, but no effect of GDM with insulin treatment relative to either lean women or women with obesity was observed (Figure 1C). By contrast, no effect of maternal obesity on syncytin-2 transcript levels was detected, however a significant ∼2.3-fold increase with GDM and insulin treatment was seen relative to women with maternal obesity (Figure 1C).

### Identification of PEG3 DNA elements and the effect of maternal obesity with or without GDM and insulin treatment on PEG3 binding

The genomic sequences containing the hPL LCR and the five hPL/GH genes were analyzed using a consensus eight nucleotide DNA element for PEG3 (5′-gkGGswsT-3′) (Lee et al. 2015). Possible high affinity PEG3 binding sites were identified in placenta-specific HS IV as well as HS III and HS V-related sequences (Figure 2A). A highly related site was also detected at the equivalent location about 0.5 kb upstream of the transcription initiation sites of all five hPL/GH genes as described (Cattini et al. 2020). EMSA with a radiolabelled high affinity PEG3 DNA element from the Pgm2L1 gene (Thiaville et al. 2013) and JEG-3 nuclear extract, was used to assess each of the sequences identified in the LCR and hPL/GH-V promoter regions for PEG3 binding through competition (Figure 2B). Three complexes of low, medium and high mobility were detected when the Pgm2L1 oligonucleotide was used as a radiolabelled probe with JEG-3 nuclear protein (Figure 2B). The low and high mobility complexes were assigned as monomer and dimer complexes with high affinity/specificity based on efficient competition with excess unlabeled Pgm2L1 oligonucleotide *versus* that seen with an oligonucleotide containing the distal *hPL-A* promoter sequences with a disrupted GG-T motif (mPL-Ap; Figure 2B). The low and high mobility PEG-3 complexes were competed with increasing HS III-V-related oligonucleotides and also, although with a lower affinity, oligonucleotides containing the distal *hPL-A* promoter region with the conserved putative PEG3-like site (Figure 2B).

**Figure 2.**
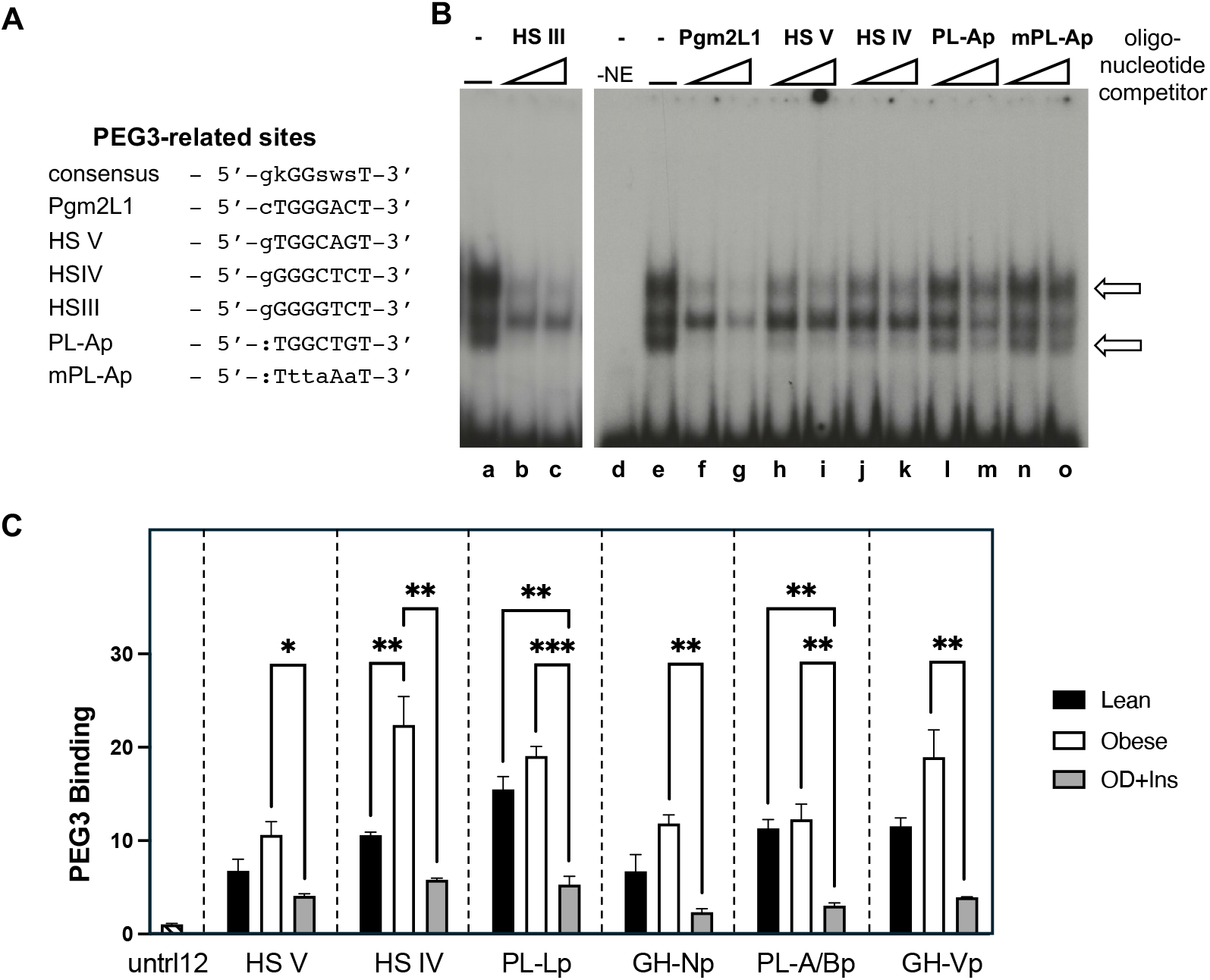
**(A)** PEG3-related DNA binding sites, including consensus and high-affinity Pgm2L1 sites, alongside hGH/PL-related sequences. (GG-T motif; k (keto) = G or T; s (strong) = G or C; and w (weak) = A or T). **(B)** Competition-EMSA with JEG-3 nuclear extract (NE) and radiolabeled Pgm2L1 probe competed with hPL gene sequences, visualized by autoradiography. Gel lanes correspond to (a and e) Pgm2L1 probe and NE, (d) Pgm2L1 probe alone, as well as Pgm2L1 probe and NE in combination with a 50 and 100 pM excess of the following oligonucleotide competitors: (b and c) HS III; (f and g) Pgm2L1; (h and i) HS V; (j and k) HS IV; (l and m) distal hPL-A promoter; and (n and o) modified distal hPL-A promoter (mPL-Ap). Three major bands are detected including a monomer and dimer complex (arrows) based on self-competition (lanes e-g). **(C)** ChIP-qPCR assessed PEG3 binding at the LCR and hGH/PL gene loci in term placentas from lean women (L), women with obesity (O), and with O/GDM+Ins. Binding events were calculated from immunoprecipitated/input DNA using specific PCR primers for LCR and hGH/PL gene-related sequences and a control untranscribed region of human chromosome 12 (Untr12). The results are expressed as mean binding events plus and minus standard error of the mean relative to the background (Ctrl), which is arbitrarily set to 1. Significance was determined by one-way ANOVA with the Tukey’s *post hoc* test for each region assessed (n=3). Significant differences are indicated by: ^*^ p<0.05; ^**^ p<0.01 and ^***^ p<0.001.

The impact of maternal obesity on PEG3 protein association with HS V and HS IV-related sequences in the LCR as well as conserved sequences in the upstream hPL/GH promoter regions was assessed by ChIP-qPCR. Human term placental samples were analyzed from three groups of pregnant women: lean (L), with obesity (O), and with O/GDM+Ins. PEG3 binding to the positive control genes *MYOF, TEF-1* and *GLTSCR1* above background Untr12 levels was confirmed in lean term placenta (data not shown). An increase in PEG3 association with the LCR was seen with maternal obesity, including a significant ∼2.1-fold (p=0.0078, n=3) increase at placenta-specific HS IV-related sequences (Figure 2C). However, binding to HS IV and HS V-related sequences was reduced significantly 74% (p=0.0014, n=3) and 63% (p<0.0133, n=3), respectively in women with O/GDM+Ins when compared to obesity alone (Figure 2C). A pattern of modest ∼1.5-fold increases in PEG3 binding to hPL/GH promoter-related sequences was suggested in placenta samples from group O *versus* L, but did not reach significance (Figure 2C). However, significant 72-80% (p<0.005, n=3) decreases in PEG3 binding to all promoter regions was detected when term placenta samples from groups O/GDM+Ins and O were compared (Figure 2C).

### Effect of maternal obesity with or without GDM and insulin treatment on placental LCR (HS III) interactions with P sequences and 3′-RRs as well as between P sequences and 3′-RRs

A relatively higher ligation frequency level was seen between HS III and the *hPL-L* P sequences (86.6%) *versus* HS III with either *hPL-A* (12.3%) or *hGH-V* and *hPL-B* P sequences combined (<1.5%) in placentas from lean women (Figure 3). However, maternal obesity was associated with a reduction in the dominance of the HS III and *hPL-L* P sequences interaction (down to 41.35%). This was accompanied by induction of an interaction between HS III and *hPL-B* P sequences (28.55%) as well as an increase in the detection of HS III and *hGH-V* interactions (23.6%), all consistent with a change in the architectural organization containing the LCR and hGH/PL genes. This pattern was also seen in women with O/GDM+Ins, except for a decrease in the HS III and *hGH-V* P sequence interaction (Figure 3).

**Figure 3.**
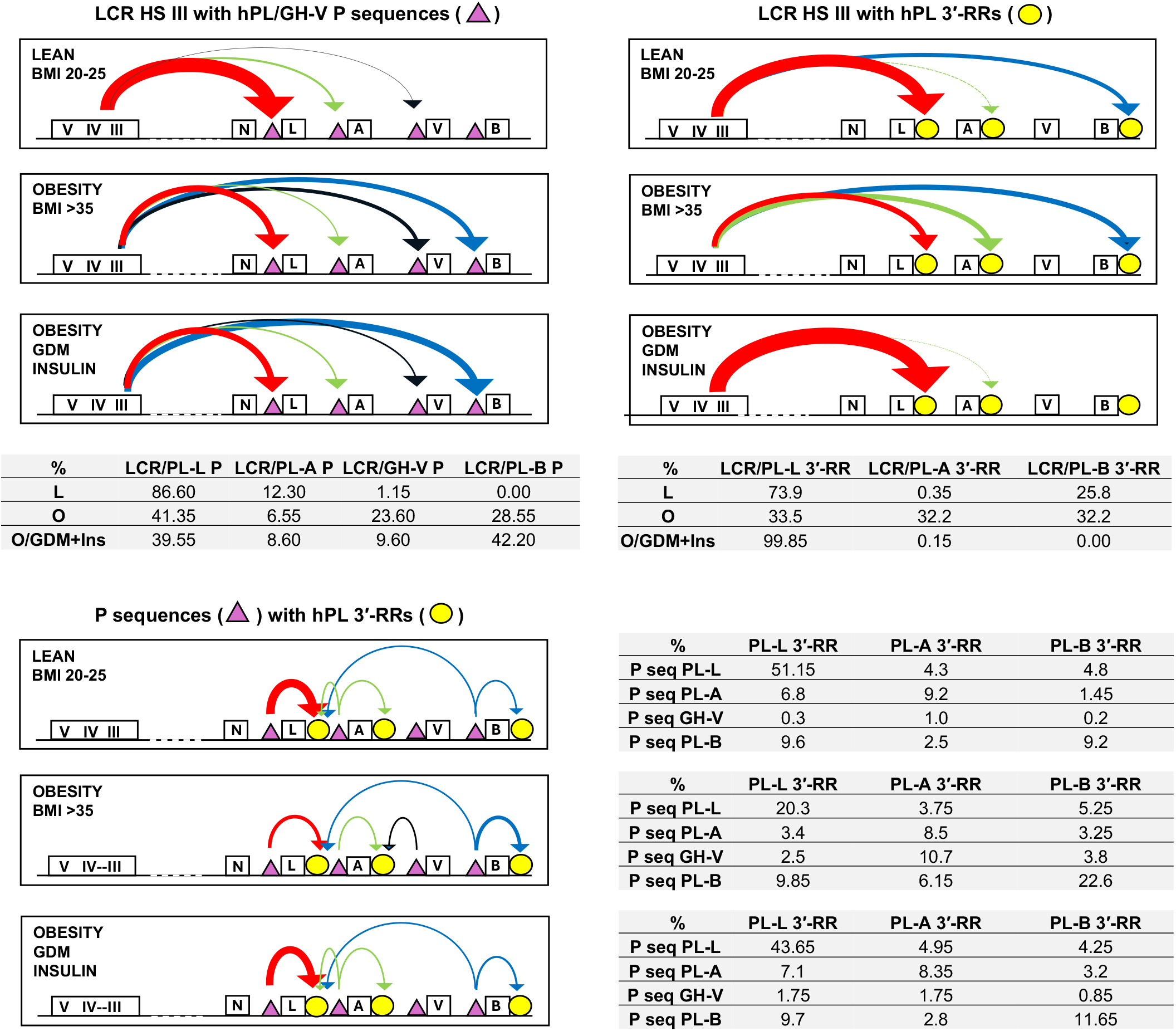
3C assay evaluated HS III interaction with P sequences (P seq) and hPL downstream regulatory regions (3′-RRs) in pooled placenta samples (n=7-9) from three groups of pregnant women: lean (L); with obesity (O); and with O/GDM+Ins. Values are expressed as a percentage and normalized for GAPDH DNA. The total interactions for L were arbitrarily set to 100% to allow comparisons and values for percentage of all interactions assessed are tabulated. The tabulated results are also presented graphically to show the major ligations detected, and arrow thickness gives an approximation of relative frequency of interaction.

A relatively higher ligation frequency level was also seen between HS III and the *hPL-L* 3′-RR (73.9%) *versus* HS III with *hPL-B* (25.8%) and *hPL-A* 3′-RRs (<1%) in placentas from lean women (Figure 3). Again, maternal obesity was associated with a loss of dominance of the *hPL-L* 3′-RR interaction and detection of comparable interactions between HS III and 3′-RRs of *hPL-L* (33.5%), *hPL-A* (32.2%) and *hPL-B* (32.2%) (Figure 3). By contrast to the HS III and *hPL-L* P sequence interaction, the dominant HS III and *hPL-L* 3′-RR interaction seen in the placentas of lean women, was recovered and even enhanced in women with O/GDM+Ins (99.85% *versus* 74.9% ligation level). Like lean women, the frequency of ligation between HS III and *hPL-A* 3′-RR was <1%, and even the level of interaction between HS III and *hPL-B* 3′-RR was reduced (<1%) (Figure 3).

A pattern of interaction between each hPL gene’s associated upstream P sequences and downstream 3′-RRs was detected in term placenta samples from lean women (Figure 3). For *hGH-V*, which has local upstream P sequence but no downstream 3′-RR, no similar level of interaction was observed. However, the frequency of ligation of *hPL-A* and *hPL-B* P sequences with the 3′-RR for *hPL-L* as well as its own (*hPL-B* 3′-RR) were comparable (Figure 3), consistent with a close association of the three hPL genes. The pattern of interactions was similar in term placenta samples from women with obesity except there was a reduction in the frequency of ligation between *hPL-L* P sequences and *hPL-L* 3′-RR and an increase *hPL-*B P sequences with the *hPL-B* 3′-RR (Figure 3). In addition, the frequency of interaction between *hPL-A* P sequences and *hPL-L* 3′-RR was reduced, and an interaction between the *hGH-V* P sequences and the 3′-RR of the upstream and adjacent *hPL-A* was now evident (Figure 3). These changes suggest a less structured arrangement of the hPL/hGH-V genes. However, the pattern of hPL/GH-V P sequences and hPL 3′-RR interactions detected in women with O/GDM+Ins, was the same as observed in lean women (Figure 3). This suggests potential rescue of the chromosomal architecture containing the hPL/GH-V genes.

## Discussion

Using a combination of protein-DNA binding studies *in vitro* (EMSA) and *in situ* (3C assay) together with assessment of gene expression in human placental cells, including with sh-RNA knockdown, a potential role for the transcription factor PEG3 in the regulation of the hPL gene expression was pursued. PEG3 RNA was detected in both CTB-like JEG-3 and BeWo cells, with significantly higher levels observed in JEG-3 cells. This correlates with the limited fusion and lower hPL expression in these cells (Ganguly et al. 2015; Li, et al. 2023; Nickel and Cattini 1991). The presence of PEG3 binding sites in the vicinity of LCR HS IV and HS V sequences and ∼0.5 kb upstream of all five hPL/GH genes (Cattini et al. 2020), was supported by EMSA and competition with a high affinity Pgm2L1 PEG3 DNA element (Thiaville et al. 2013). Association of PEG3 with HS III-V and promoter sequences was also detected in term placenta tissue *in situ* by ChIP assay. This placental tissue is expected to include villous CTBs, the site of PEG3 production, as well as STBs where the hPL genes are efficiently expressed (Hiby et al. 2001; Liu et al. 2018).

To assess the effect of PEG3 on hPL/hGH-V gene expression, shRNA knockdown was done in JEG-3 cells, resulting in an ∼2-fold increase in hPL-A RNA levels. This suggests a loss of repression via the identified PEG3 site in distal promoter sequences. However, no equivalent effect on *hPL-B* or *hGH-V* transcript levels was seen, in spite of the presence of an equivalent PEG3 binding site (Cattini et al. 2020). This may reflect incomplete PEG3 knockdown combined with dominant expression of hPL-A transcripts, which account for over 80% of hGH/PL RNA (Jin et al. 2018). Another possibility is that the knockdown of PEG3 mimics a normal event during hPL gene locus activation, where PEG3 binding decreases as CTBs transition to STBs. The activation of the hPL locus occurs in CTBs and is characterized by nuclease sensitivity and histone modifications (Kimura et al. 2007; Nickel and Cattini 1996). Acetylation of histones H3 and H4 was noted at HS III and HS V during the CTB to STB transition, with significant acetylation observed at HS IV and across the hPL locus in term STBs (Kimura et al. 2007). This is consistent with PEG3’s role in repressing mouse PL2 gene expression in spongiotrophoblast cells (Kim, et al. 2013; Tunster, et al. 2018), which develop into trophoblast giant cells, the functional equivalent to STBs in humans, that express PL2 as a key product during the second half of murine pregnancy (Simmons and Cross 2005; Soares 2004; Soncin, et al. 2015).

The hPL genes were among the first cloned (Fiddes, et al. 1979; Shine, et al. 1977), however, the factors regulating the activation and expression of hPL genes in villous STBs remain poorly defined. Local regulatory regions, including the 3′-RR, have been implicated in de-repressed and/or enhanced hPL promoter activity (Chen et al. 1989; Elefant et al. 2000b; Jacquemin et al. 1994; Jiang et al. 1999; Lytras and Cattini 1994; Lytras et al. 2011; Norquay et al. 2006; Vakili et al. 2013; Walker et al. 1990). This includes CCAAT-enhancer-binding protein (C/EBP) β, which in contrast to PEG3 is produced at higher levels in STBs compared to CTBs (Bamberger, et al. 2004), and decreased C/EBPβ binding is associated with reduced gene expression (Vakili et al. 2013). However, consistent placenta-specific expression of the hPL genes is only seen in transgenic mice when sequences containing HS III-V, in addition to local regulatory and promoter sequences are included (Cattini et al. 2020; Jin, et al. 2009; Jones et al. 1995).

In a prior 3C study of the hGH/PL gene locus in placental cells, HS III-V sequences were suggested to interact with *hGH-N* and *hGH-V* promoters but not with hPL genes (Tsai et al. 2016). A schematic representation of these interactions is reproduced here for comparison with the interaction detected by us, which includes the hPL genes (Figure 4). Specifically, when interactions with LCR HS III-V were examined using a *Bgl* II *versus Sac* I fragments and a HS III *versus* HS V anchor primer, interactions with all three hPL gene-related sequences were confirmed (Jin et al. 2018). This supports the idea that the LCR contributes to establishing the relatively inactive hGH and active hPL gene domains in term placenta (Figure 4). HS V sequences interact with *hGH-N* and *hGH-V*, serving as an upstream boundary for the hGH gene domain, which has a less open chromatin structure (Jin et al. 2011; Nickel and Cattini 1996). Conversely, HS IV binds CTCF and acts as a placenta-specific boundary element for the hPL gene domain, which exhibits a more open chromatin structure conducive to transcriptional activity (Kimura et al. 2007; Nickel and Cattini 1996). The hPL gene domain relies on interactions between HS III-related sequences and upstream *hPL-L*, which may facilitate interactions with *hPL-A* and *hPL-B* via the 3′-RR and P sequences. This structural organization positions HS III and enhancer activity in the proximity of hPL genes (Jin et al. 2004), which may explain higher expression levels in term placenta (Jin et al. 2018; Vakili et al. 2013). Conversely, HS IV sequences may limit enhancer activity on the hGH gene domain. The minimal expression of *hGH-V* (Ganguly et al. 2015; Tsai et al. 2016), may reflect its location between highly expressed *hPL-A* and *hPL-B* (Kimura et al. 2007; Nickel and Cattini 1996), but a role for hGH-V P sequences cannot be excluded.

**Figure 4.**
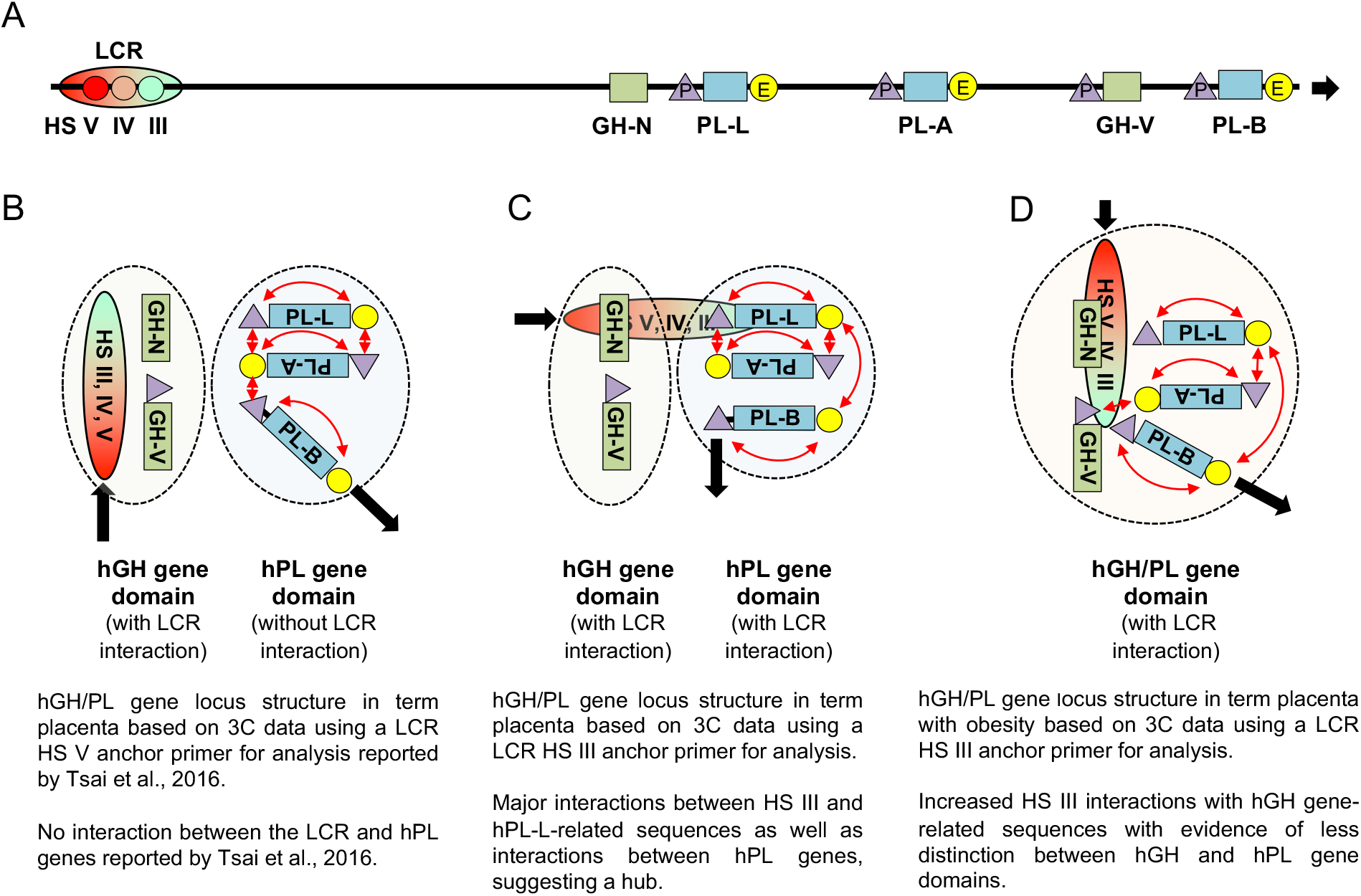
Schematic summarizing the human (h) GH/PL gene domain interactions in term placenta samples from women with and without maternal obesity by 3C assay. (**A**) Components of the GH/PL gene locus include GH-N/GH1, PL-L/CSHL1, PL-A/CSH1, GH-V/GH2 and PL-B/CSH2 genes, hypersensitive site (HS) III, IV and V sequences which make up the LCR in the placenta, local upstream P (triangle) and downstream regulatory region enhancer (E) sequences (circle). GH and PL gene domain interactions detected in term placenta using a **(B)** HS V as reported in reference (Tsai et al. 2016) or **(C)** HS III anchor primer as reported here. (**D**) GH and PL gene interactions detected in women with maternal obesity using a HS III anchor primer. Interactions between PL P and E sequences and between PL genes are indicated (red arrows).

Maternal obesity disrupts this chromosomal architecture and hPL gene expression (Jin et al. 2018; Vakili et al. 2013), resulting in a loss of distinction between hGH and hPL gene domains (Figure 4). This correlates with sustained or increased PEG3 binding at HS IV in term placentas, which would interfere with any requirement for decreased CTB PEG3 binding that may potentially facilitate hPL gene locus activation and efficient expression in STBs. The disrupted locus structure with maternal obesity is further supported by observed reductions in the interactions between HS III and hPL gene sequences, and new interactions with *hGH-N* and *hGH-V*-related sequences. Also, the favored interaction between HS III and *hPL-L* is lost as similar levels of interaction are now seen with *hPL-A, hPL-B* and *hPL-L* sequences, and loosening of the arrangement of genes between and/or within domains is reflected in the pattern of P sequences and 3′-RR interactions.

It is not possible to distinguish whether these different interactions observed in term placenta samples occur in a single nucleus or are the combination of individual interactions from multiple nuclei. However, in cases of maternal obesity, structural changes in the locus may be reversible during pregnancy. If these changes occur prior to GDM diagnosis, insulin treatment can partially restore the two gene domains, resulting in increased hPL/hGH-V gene expression (Jin et al. 2018). This includes re-establishing interactions between HS III and *hPL-L* 3′-RR sequences, along with a rescue of the close association of hPL genes, which aligns with findings in placentas from lean women. Furthermore, this rescue of the hPL gene domain is associated with decreases in PEG3 binding across the LCR and hGH/PL sequences. Insulin treatment may target PEG3 binding generally based on similar negative effects with MYOF, TEF-1 and GLTSCR1 gene sequences (data not shown).

Decreases in PEG3 and hPL RNA levels (>40%) have been linked to an increased risk for peripartum depression (PPD) (Janssen, et al. 2016). It is unlikely that this reflects a direct link between decreases in PEG3 availability and hPL gene transcription, given the effects of PEG3 knockdown observed and the different trophoblast sites for PEG3 and hPL production, but rather a negative effect on placenta and trophoblast development (Cattini et al. 2020; Hiby et al. 2001; Liu et al. 2018; Mayhew 2014; McWilliams and Boime 1980). In this context, maternal obesity has been linked to placenta and trophoblast disruptions during pregnancy (Brouwers, et al. 2019; Leon-Garcia, et al. 2016; Turowski and Vogel 2018) and an increased risk of PPD (Jarmasz, et al. 2021; Molyneaux, et al. 2014; Steinig, et al. 2017). However, there was no correlation between PEG3 RNA levels and the pattern of PEG3 binding to HS IV sequences seen in term placenta samples from women with maternal obesity or O/GDM+Ins.

Syncytins-1 and 2 are expressed by villous trophoblasts and play essential roles in the CTB transition to STBs. Syncytin-1 positively affects both maintenance of a PEG3-expressing CTB pool for fusion throughout pregnancy and CTB fusion (Soygur and Sati 2016), while syncytin-2 plays a more important role in fusion alone (Priscakova, et al. 2023; Vargas et al. 2009). Maternal obesity was associated with a decrease in syncytin-1 RNA levels. Thus, any change in the proportion of a particular PEG3-expressing CTB subtype related to a decrease in proliferation and/or fusion with maternal obesity may not be detected. However, increased PEG3 binding to the LCR may still be observed as locus activation starting in the CTB is delayed due to modifications or additional factors required prior to fusion (Tsai et al. 2016). By contrast, syncytin-2 RNA levels were increased in women with O/GDM+Ins. This correlates with decreased PEG3 binding to the hPL LCR and other sequences with insulin treatment as well as the increased hPL gene expression and suggested decrease in the risk of PPD observed (Jarmasz et al. 2021; Jin et al. 2018).

In summary, PEG3 binds sequences in the hPL LCR, including HS IV, with high affinity. A reduction in PEG3 binding is associated with hPL locus activation and elevated *hPL-A* expression. The organization of the hGH/PL genes into two distinct hGH and hPL gene domains, with the hPL gene domain exhibiting a more open structure, highlights the regulatory mechanisms at play. Maternal obesity is associated with continuing or increased PEG3 binding at HS IV and disruption of this domain architecture and hPL gene expression, while insulin treatment (for GDM) can partially restore the gene domain structure and rescue hPL/GH-V production.

## Acknowledgements

This work was supported by a grant from the Canadian Institutes of Health Research (FRN-166215). PAC holds the H.G. Friesen Chair in Endocrine & Metabolic Disorders.

## Notes

Conflicts of Interest: The authors of the manuscript have no conflicts of interest to declare.

### Competing Interest Statement

The authors have declared no competing interest.

